# The highly conserved stem-loop II motif is important for the lifecycle of astroviruses but dispensable for SARS-CoV-2

**DOI:** 10.1101/2022.04.30.486882

**Authors:** Andrew B Janowski, Hongbing Jiang, Chika Fujii, Macee C Owen, Traci L Bricker, Tamarand L Darling, Houda H. Harastani, Kuljeet Seehra, Stephen Tahan, Ana Jung, Binita Febles, Joshua A Blatter, Scott A Handley, Bijal A Parikh, Valeria Lulla, Adrianus CM Boon, David Wang

## Abstract

The stem-loop II motif (s2m) is an RNA element present in viruses from divergent viral families, including astroviruses and coronaviruses, but its functional significance is unknown. We created deletions or substitutions of the s2m in astrovirus VA1 (VA1), classic human astrovirus 1 (HAstV1) and severe acute respiratory syndrome coronavirus 2 (SARS-CoV-2). For VA1, recombinant virus could not be rescued upon partial deletion of the s2m or substitutions of G-C base pairs. Compensatory substitutions that restored the G-C base-pair enabled recovery of VA1. For HAstV1, a partial deletion of the s2m resulted in decreased viral titers compared to wild-type virus, and reduced activity in a replicon system. In contrast, deletion or mutation of the SARS-CoV-2 s2m had no effect on the ability to rescue the virus, growth *in vitro*, or growth in Syrian hamsters. Our study demonstrates the importance of the s2m is virus-dependent.

## Background

In 1997, Monceyron *et al* described a secondary structure present near the 3′ end of classic human astroviruses named the stem-loop II motif (s2m), and a similar element was also identified in avian bronchitis virus, a coronavirus^1^. Subsequently, s2m elements have been detected in members of the *Astroviridae, Caliciviridae, Coronaviridae, Picornaviridae*, and *Reoviridae* viral families, all with highly conserved nucleotide sequences of 39-43 nucleotides in length (**Fig. 1A-E**)^2–6^. Currently, the function or necessity of the s2m for the viral lifecycle is poorly understood. The phylogenetic distribution suggests horizontal acquisition of the s2m at different timepoints, and maintenance of the element suggests that it confers a fitness advantage^2, 3^. The crystal structure of the s2m from severe acute respiratory syndrome coronavirus 1 (SARS-CoV-1) demonstrates a tertiary structure that includes a 90° kink in the helix axis that results in additional tertiary interactions^4^. Antisense oligonucleotides to the s2m reduced viral replication for severe acute respiratory syndrome coronavirus 2 (SARS-CoV-2) and classic human astrovirus 1 (HAstV1) replicons^7^. These results suggest that the secondary structure of the s2m is important for virus replication.

**Figure 1:**
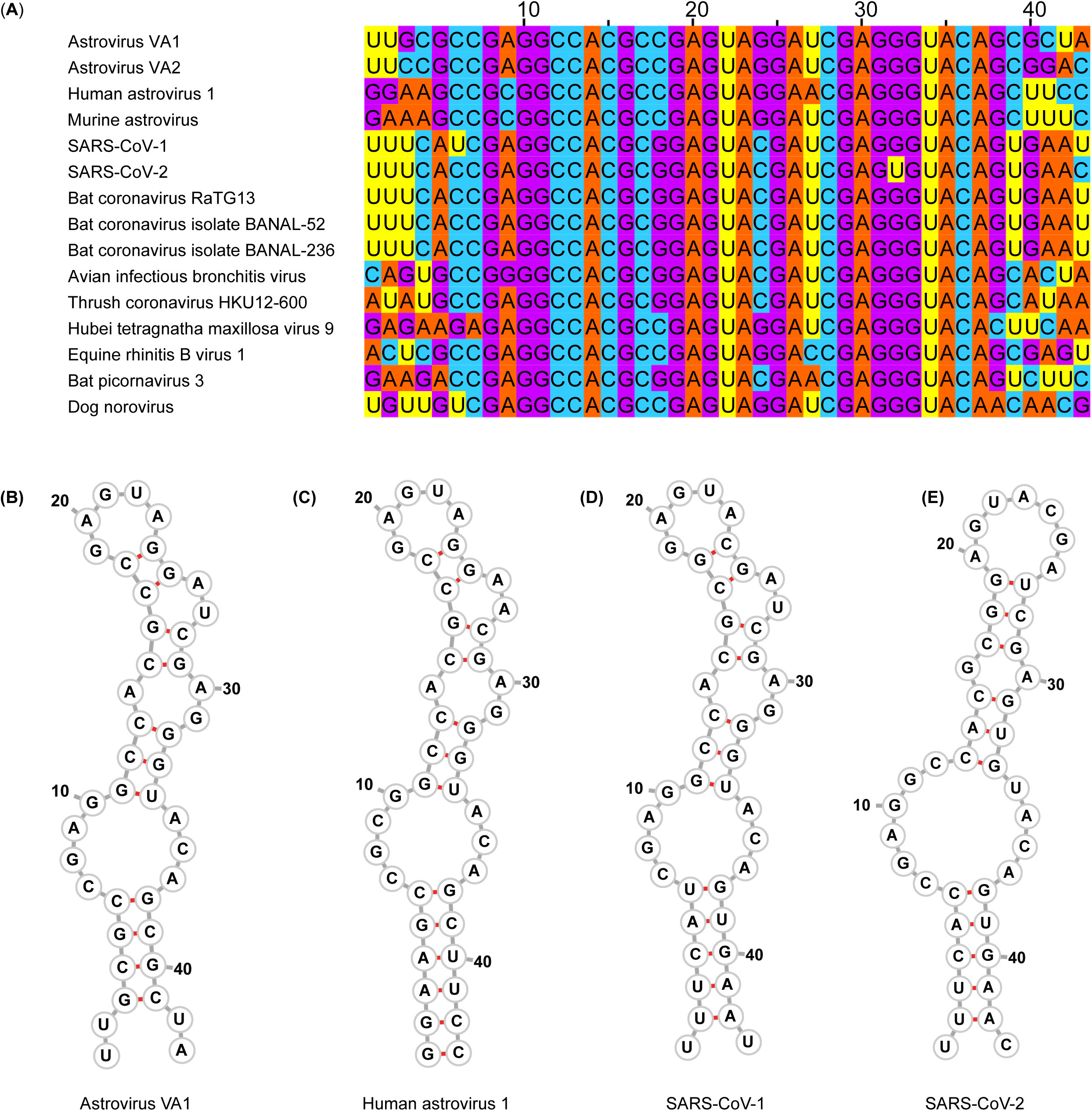
s2m nucleotide sequences and predicted secondary structures. (A) Multiple sequence alignment of representative viruses that encode an s2m. (B-E) Predicted secondary structure of the s2m by RNAfold for (B) Astrovirus VA1, (C) Human astrovirus 1, (D) SARS-CoV-1, (E) SARS-CoV-2

Astroviruses are non-enveloped, single-stranded, positive-sense RNA viruses that are approximately 6,000-7,700 nucleotides in length^8, 9^. Astroviruses are associated with infections of the gastrointestinal and nervous systems and harbor additional tissue tropisms^9^. Many astrovirus species encode an s2m near the end of open reading frame 2 (ORF2)^2^. Of the three clades of astroviruses known to infect humans (HAstV, VA, and MLB astroviruses)^8^, members of the HAstV and VA clades contain an s2m, while MLB astroviruses lack an s2m^10^. Previously established culture systems exist for astrovirus VA1 (VA1) and HAstV1^11–14^, with published reverse genetics and replicon systems for HAstV1^15, 16^.

Coronaviruses are enveloped, single-stranded, positive-sense RNA viruses that are approximately 30,000 nucleotides in length^17^. Four genera comprise the coronavirus family, with some members of the beta-(sarbecovirus subgenus), gamma-, and delta-genera containing an s2m^3^. Both SARS-CoV-1 and SARS-CoV-2 encode an s2m in the 3′ untranslated region (UTR)^3, 18^. In contrast, seasonal human coronaviruses (HKU1, 229E, OC43, and NL63) and Middle Eastern respiratory syndrome coronavirus (MERS-CoV) do not contain an s2m^3^. Interestingly, the reference SARS-CoV-2 genome and all subsequent variants of concern encode a unique genetic variant with a uracil at position 32 in the s2m that is distinct from other s2m sequences and is predicted to perturb the secondary structure^19–23^. Additional genetic variants or deletions of the s2m have been periodically detected from clinical isolates^23–26^.

We determined the functional significance of the s2m for the viral lifecycle of three viruses, VA1, HAstV1, and SARS-CoV-2 by using reverse genetics to mutagenize the s2m element as well as by studying a clinical isolate of SARS-CoV-2 that has a natural deletion in the s2m region.

## Results

### VA1 s2m is essential for viral propagation

A reverse genetics system for VA1 was developed using the reference genomic sequence (NC_013060.1; **Fig. S1A**). VA1 was consistently rescued with a median viral quantity of 9.4×10^5^ focus forming units (FFU)/mL after serial passage (**Table 1**, **Fig. 2A**). In multi-step growth curves in Caco-2 cells, viral RNA increased >1,000-fold in 24 hours with further increases over time, consistent with a published strain that has been passaged seven times in Caco-2 cells (VA1-C-P7; **Fig. S1B**)^11, 12^. We then determined the effect of deletion of the s2m in VA1. Because the predicted stop codon for ORF2 of VA1 is located within the s2m, the last 16 nucleotides of the s2m downstream of the stop codon (named for the nucleotide positions in the multiple sequence alignment: VA1-s2m^Δ25–41^; **Table S1** and **Fig. 1A**) were deleted. Compared to the wild-type (WT) virus, VA1-s2m^Δ25-41^ could not be recovered in all 9 attempts (**Table 1**, **Fig. 2A**), suggesting the s2m is critical for VA1.

**Figure 2:**
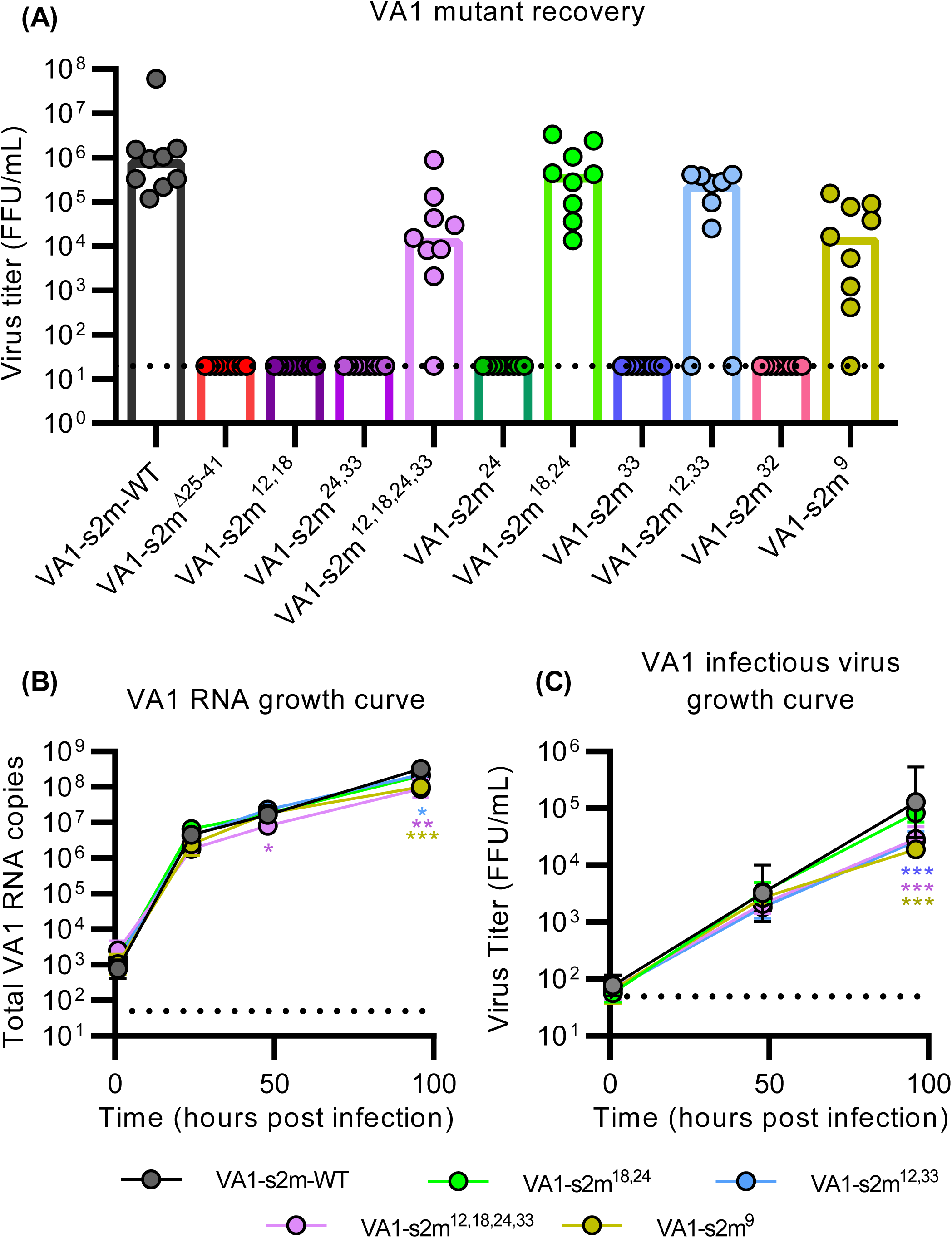
The s2m is essential for astrovirus VA1. (**A**) Nine replicates of WT or mutant VA1 genomes were transfected and serially passaged. Disruption of the VA1 s2m was achieved through deletion (s2m^Δ25–41^), and single (VA1-s2m^24^ and VA1-s2m^33^) or double substitutions (VA1-s2m^12, 18^ and VA1-s2m^24, 33^) that are predicted to significantly alter the secondary structure of the s2m. The loss of the single or double G-C binding pairs was complemented with compensatory substitutions that are predicted to restore the s2m secondary structure: VA1-s2m^18, 24^, VA1-s2m^12, 33^, and VA1-s2m^12, 18, 24, 33^. VA1-s2m^32^ contains the unique uracil genetic variant encoded by the SARS-CoV-2 s2m. VA1-s2m^9^ contains a single substitution at a position that does not form a G-C binding pair. Plotted are the individual virus titers, bars are the median value, and the dashed line is the limit of detection. (**B-C**) Multi-step growth curve of VA1 recombinant viruses that were recoverable using an MOI of 0.01 in Caco-2 cells. A total of three experiments with three technical replicates each were conducted. For both graphs, plotted are geometric means and bars representing the geometric standard deviation. The dashed line is limit of detection. ** denotes P≤ 0.01, *** denotes P≤ 0.001. (**B**) Viral RNA multi-step growth curve with a significant difference identified between WT and mutants by two-way ANOVA (F[4,40]= 14.57, P< 0.001). Compared to VA1-s2m-WT, VA1-s2m^12, 18, 24, 33^ was lower at 48 and 96 hpi (48 hpi P= 0.045, 96 hpi P= 0.0014). VA1-s2m^9^ and VA1-s2m^12, 33^ had lower RNA at 96 hpi compared to WT (VA1-s2m^9^: P< 0.001, VA1-s2m^12, 33^ P= 0.049), while VA1-s2m^18, 24^ had no difference (P> 0.06). (**C**) Multi-step growth curve of virus titers with a significant difference between WT and mutants identified by two-way ANOVA (F[4,40]= 11.4, P< 0.001). Reduced viral titers for VA1-s2m^9^, VA1-s2m^12, 33^, and VA1-s2m^12, 18, 24, 33^ were detected at 96 hpi compared to WT (all P< 0.001).

**Table 1:** Frequency of recovery of VA1 s2m mutants. . A total of 9 attempts were made to rescue each VA1 s2m recombinant virus. Presence of infectious virus was detected using a focus forming assay.

| s2m name | Frequency of recovery (%) |
| --- | --- |
| VA1-s2m-WT | 9/9 (100) |
| VA1-s2m<sup>Δ25-41</sup> | 0/9 (0) |
| VA1-s2m<sup>12,18</sup> | 0/9 (0) |
| VA1-s2m<sup>24,33</sup> | 0/9 (0) |
| VA1-s2m<sup>12,18,24,33</sup> | 8/9 (89) |
| VA1-s2m<sup>24</sup> | 0/9 (0) |
| VA1-s2m<sup>18,24</sup> | 9/9 (100) |
| VA1-s2m<sup>33</sup> | 0/9 (0) |
| VA1-s2m<sup>12,33</sup> | 7/9 (78) |
| VA1-s2m<sup>9</sup> | 8/9 (89) |
| VA1-s2m<sup>32</sup> | 0/9 (0) |

### G-C binding pairs are critical for the function of the VA1 s2m

To further corroborate the importance of the s2m, we created substitutions of specific nucleotides of the s2m (**Table S1**). We first designed a mutant that significantly altered the predicted secondary structure of the s2m by creating a double mutant that contained replacement of cytosines with guanines at positions 12 and 18 (VA1-s2m^12, 18^; **Fig. S2)**. These substitutions affect two guanine-cytosine (G-C) base pairs that are highly conserved in the multiple sequence alignment (**Fig. 1A** and **S2**). Importantly, these substitutions are synonymous and do not alter the amino acid sequence of ORF2. Attempts to rescue and grow VA1-s2m^12, 18^ were not successful in all nine attempts (**Table 1**, **Fig. 2A**). We also created substitutions at positions 24 and 33 that form G-C base pairs with positions 12 and 18 (**Table S1**; VA1-s2m^24, 33^). Both guanines were converted to cytosines, which also alters the predicted secondary structure (**Fig. S2)**. The substitution at position 24 disrupts the stop codon resulting in addition of 16 amino acids to ORF2 (**Table S2**). Like VA1-s2m^12, 18^, we could not rescue VA1-s2m^24, 33^ (**Table 1**, **Fig. 2A**). Next, we determined the impact of altering a single G-C base pair by creating point substitutions at position 24 alone (VA1-s2m^24^) and position 33 alone (VA1-s2m^33^; **Table S1**). In both mutants, the guanine was converted to a cytosine, resulting in slightly altered s2m secondary structures (**Fig. S2**). VA1-s2m^24^ results in disruption of the stop codon and adds 16 amino acids to ORF2 while VA1-s2m^33^ does not affect the amino acid sequence (**Table S2**). No virus could be propagated from either VA1-s2m^24^ or VA1-s2m^33^ in all 9 rescue attempts, respectively (**Table 1, Fig. 2A**). These results demonstrate that disruption of even a single G-C base pair impacts the biological activity of the VA1 s2m.

Next, we determined if the function of the mutant s2m sequences could be restored by creating complementary substitutions that would reestablish G-C binding pairs but invert them relative to the WT s2m. Using the VA1-s2m^33^ mutant as a template, we converted position 12 from a cytosine to guanine (VA1-s2m^12, 33^), which is a synonymous substitution (**Table S1-S2**). For VA1-s2m^24^, we altered position 18 by also converting the cytosine to guanine (VA1-s2m^18, 24^), also a synonymous substitution (**Table S1-S2**). This mutant maintains the altered nucleotide at the stop codon resulting in addition of 16 amino acids (**Table S2**). The secondary structures for both VA1-s2m^12, 33^ and VA1-s2m^18, 24^ are predicted to form the WT s2m structure (**Fig. S2)**. Both reversion mutants yielded viable virus in 7 out of 9 replicates for VA1-s2m^12, 33^ and 9 out of 9 replicates for VA1-s2m^18, 24^ (**Table 1**, **Fig. 2A**). Because VA1-s2m^18, 24^ was viable despite alteration of the stop codon, this finding suggests that the extension of ORF2 by 16 amino acids does not by itself compromise the viability of VA1. Finally, we determined if we could recover a mutant in which two G-C base pair sites were inverted compared to WT. We created a quadruple mutant s2m (VA1-s2m^12, 18, 24, 33^), that combines the substitutions of VA1-s2m^12, 18^ and VA1-s2m^24, 33^ that individually were unrecoverable (**Table S1**). VA1-s2m^12, 18, 24, 33^ is predicted to form the same secondary structure as the WT s2m and contains the substitution that disrupts the stop codon (**Fig. S2 and Table S2**). VA1-s2m^12, 18, 24, 33^ was recovered in 8 out of 9 replicates (**Table 1**, **Fig. 2A**), further confirming the importance of the G-C base pairs in the s2m and that VA1 can tolerate two G-C inversions.

The SARS-CoV-2 s2m encodes uracil at position 32 while essentially all other s2m sequences known to date contain guanine (**Fig. 1A**). We further analyzed 295,000 complete SARS-CoV-2 genomes uploaded to the NCBI database from 2020-2021. Of those sequences, there were only 4 genomes with guanine, 5 with cytosine, and 6 with small deletions while all other sequences contained uracil at position 32. In the canonical s2m secondary structure, this position is predicted to form a G-C base pair, and based on our previous results, we hypothesized that VA1 will not tolerate this substitution. To determine if VA1 could tolerate a uracil substitution at this position, we created VA1-s2m^32^ (**Table S1** and **Fig. S2**). Similar to the other mutant VA1 s2m sequences with disrupted G-C binding pars, VA1-s2m^32^ was not rescued in nine attempts (**Table 1**, **Fig. 2A**).

We also noticed in the multiple sequence alignment that position 9 can be variable between viruses (**Fig. 1A**), including viruses within the same family. VA1, SARS-CoV-1, and SARS-CoV-2 encode an adenine while classic human astroviruses encode a cytosine. This position is not predicted to form a G-C base pair (**Fig. 1B)**, but has been identified to form potential long-distance tertiary interactions with nucleotide 30 in the SARS-CoV-1 s2m crystal structure^4^. To evaluate the significance of this position and nucleotide residue, we mutated position 9 in the VA1 s2m by replacing the adenine for a cytosine (VA1-s2m^9^; **Table S1**). The VA1-s2m^9^ mutant could be propagated in 8 out of 9 replicates (**Table 1**).

Lastly, we determined if any of the recoverable VA1 mutants had a growth defect relative to WT. For all mutants that were propagated, we confirmed the presence of the expected s2m sequence with the engineered substitutions by Sanger sequencing. We selected one isolate from VA1-s2m-WT, VA1-s2m^9^, VA1-s2m^18, 24^, VA1-s2m^12, 33^, VA1-s2m^12, 18, 24, 33^ for deep sequencing. For each recombinant virus, we obtained consensus sequences from high confidence reads that covered ≥99.7% of the genome. The expected substitutions in the s2m were identified and no other single nucleotide polymorphisms were detected in the genome. In multi-step growth curves in Caco-2 cells, there was statistically significant differences in total RNA quantity per well (two-way ANOVA F[4,40]= 14.57, P< 0.001; **Fig. 2B**) and virus titers (two-way ANOVA F[4,40]= 11.4, P< 0.001; **Fig. 2C**). Of the four mutants at 48 hours post-infection (hpi), only VA1-s2m^12, 18, 24, 33^ had lower viral RNA compared to WT (2-fold, P= 0.045), but there was no difference in the viral titer for any mutant (all P> 0.8). At 96 hpi, viral RNA and titers were reduced for VA1-s2m^12, 18, 24, 33^ (RNA: 3.6-fold, P= 0.0014; virus titer: 4.4-fold, P< 0.001), VA1-s2m^12, 33^ (RNA: 1.5-fold, P= 0.049; virus titer: 4.9-fold P< 0.001) and VA1-s2m^9^ (RNA: 3.3-fold, P< 0.001; virus titer: 6.8-fold, P< 0.001). VA1-s2m^18, 24^ had no significant differences in RNA or viral titer (both P> 0.06). These results suggest that some of the recombinant viruses have a growth defect, with VA1-s2m^9^ having the greatest reduction at 96 hpi.

### The s2m is important for classical human astrovirus 1

To demonstrate the functional significance of the s2m in a different astrovirus, we introduced a partial deletion of the s2m in HAstV1 (**Table S1**) using a published reverse genetics system^15^. As with the VA1 s2m, the HAstV1 s2m contains a stop codon for ORF2, so we introduced a deletion of 19 nucleotides after the stop codon (HAstV1-s2m^Δ25–43^). In contrast to our results with VA1, recombinant HAstV1 virus containing the deletion could be rescued and had no unexpected mutations identified by Sanger sequencing (**Fig. 3A**). We next determined if there was a growth defect of the s2m deletion mutant. In a multistep growth curve in Caco-2 cells, the HAstV1-s2m^Δ25-43^ mutant had reduced production of virus progeny compared to WT (two-way ANOVA F[1,4]= 180, P< 0.001), with 2.9- to 6-fold decreases occurring at 24 (P= 0.002), 72 (P= 0.009) and 96 (P= 0.04) hpi (**Fig. 3B**). We further corroborated this result through a HAstV1 based replicon system. In HEK293T cells, the HAstV1-s2m^Δ25-43^ replicon mutant had activity that was reduced by 80% compared to WT (P= 0.001), while in Huh7.5 cells, the HAstV1-s2m^Δ25-43^ replicon activity was reduced by 40% (P= 0.002; **Fig. 3C-D**). These results demonstrate the importance of the s2m for replication of HAstV1.

**Figure 3:**
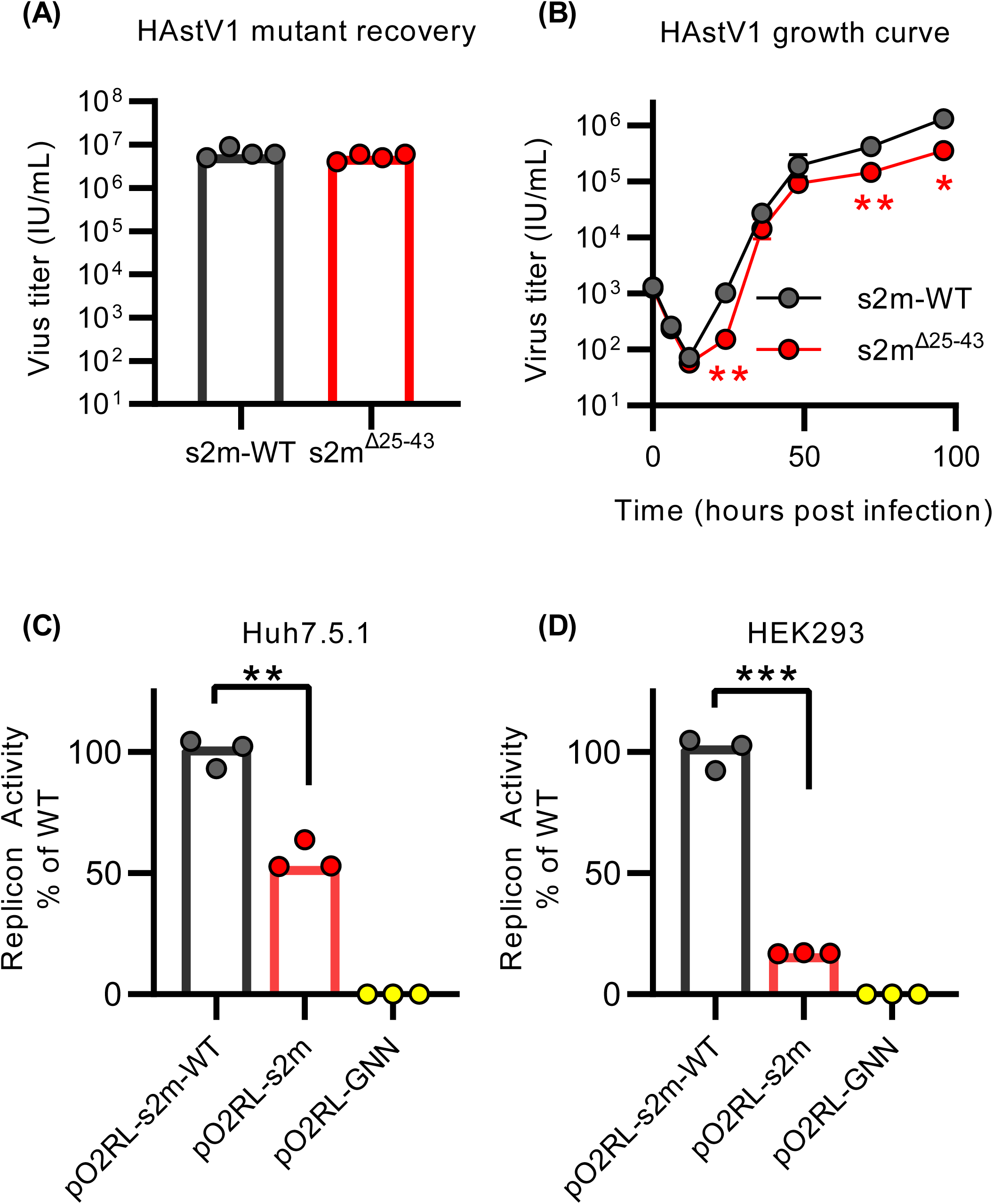
The s2m is important for human astrovirus 1. (**A**) HAstV1 genomes containing s2m-WT or a deletion of the s2m were created in quadruplicate, passaged, and viral titers were measured. (**B**) A multi-step HAstV1 growth curve was obtained by infecting Caco-2 cells with HAstV1 and HAstV1-s2m^Δ25-43^ at an MOI of 0.1. A total of three experiments were conducted. Plotted are the geometric means with error bars representing geometric standard deviations of the virus titer infectious units per mL (IU/mL). A two-way ANOVA identified reduced viral titers for HAstV1-s2m^Δ25-43^ compared to WT (F[1,4]= 180, P< 0.001), with significantly decreased titers detected at 24 (P= 0.002), 72 (P= 0.009) and 96 (P= 0.04) hours post-infection by post-hoc testing. (**C-D**) Median relative replicon luciferase activities measured after RNA transfection of (**C**) Huh7.5.1 and (**D**) HEK293T cells. Values are normalized so that the median HAstV1 replicon WT (pO2RL-s2m-WT) value is 100%, GNN mutant is RdRp knockout replicon and controls for background activity of the assay. A total of three experiments were conducted. WT and deletion replicons compared using a two-tailed T-test. For all graphs, * denotes P≤ 0.05, ** denotes P≤0.01. *** denotes P≤ 0.001

### The s2m element is dispensable for SARS-CoV-2 *in vitro*

We next determined the importance of the s2m in the context of a different viral family using SARS-CoV-2. We engineered three recombinant viruses with mutations or deletions in the s2m (**Fig. S1C)**. First, we deleted nucleotides 2-42 of the SARS-CoV-2 s2m (CoV-2-s2m^Δ2-42^) as the s2m is completely encoded in the 3′ UTR (**Table S1**). We also created a mutant that contained four consecutive nucleotide substitutions in the stem region of the s2m (CoV-2-s2m^2-5^) that is predicted to disrupt the secondary structure as well as a revertant mutant (CoV-2-s2m^2-5, 39–42^) that contained four additional compensatory substitutions that restored the predicted stem region and the SARS-CoV-2 s2m secondary structure (**Table S1**, **Fig. S3**). All mutant SARS-CoV-2 genomes yielded infectious viral particles that could be propagated in Vero-hTMPRSS2 cells. We confirmed the presence of the engineered mutations and the absence of spontaneous mutations in the SARS-CoV-2 s2m by sequencing. We next tested whether there were any defects in the growth rate of the mutants by conducting multi-step growth curves (**Fig. 4A**). There was no significant difference between the WT and mutant viruses at any timepoint in Vero-hTMPRSS2 (African green monkey) cells (two-way ANOVA F[3,20]= 1.02, P= 0.4) or Calu-3 (human) cells (two-way ANOVA F[3,20]= 0.48, P= 0.7), suggesting that the s2m element is not required for SARS-CoV-2 *in vitro* (**Fig. 4B**).

**Figure 4:**
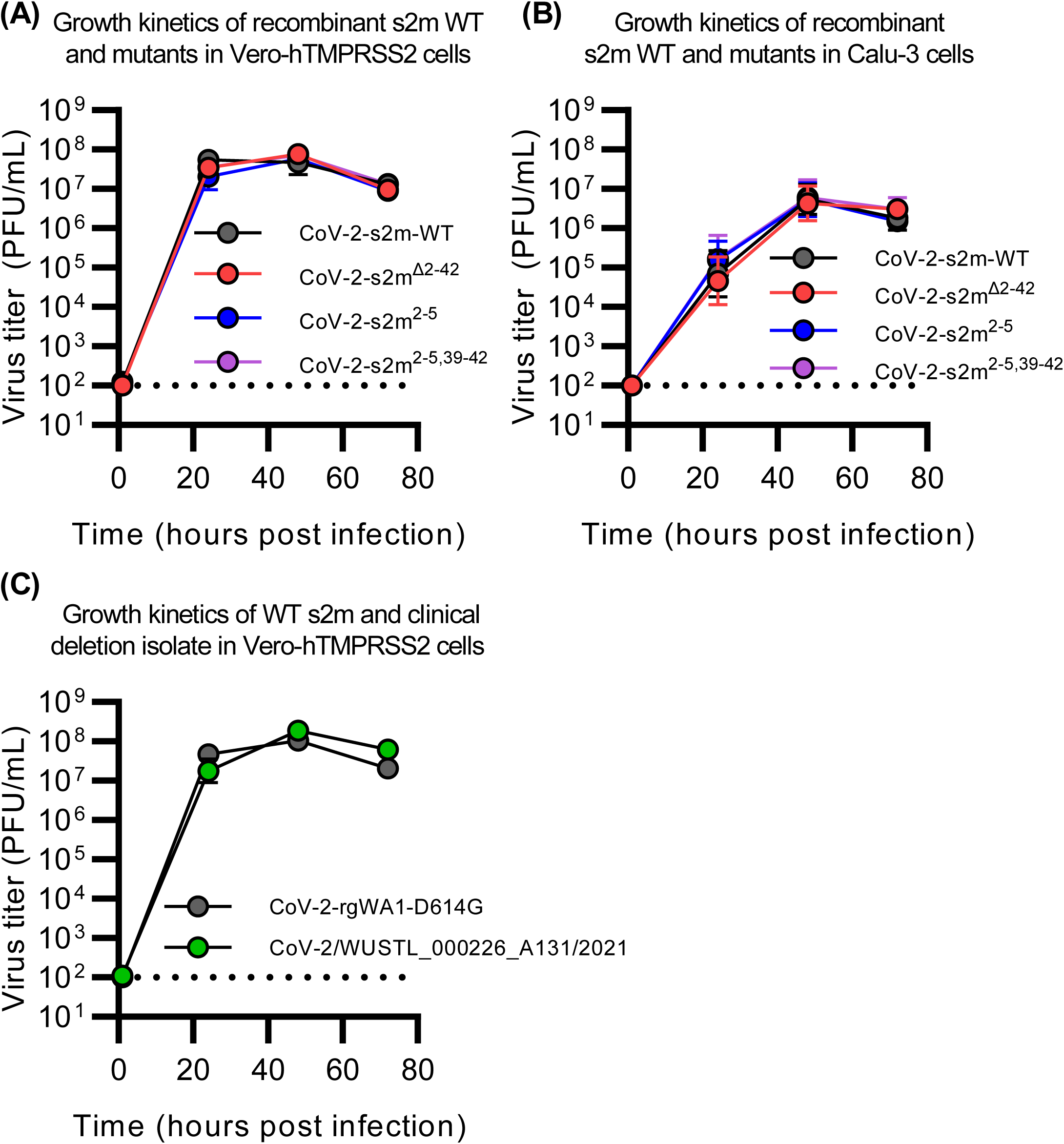
The s2m is dispensable for SARS-CoV-2 *in vitro.* (**A-B**) Multi-step growth curve of s2m-WT and mutants in (**A**) Vero-hTMPRSS2 and (**B**) Calu-3 cells. Mutant strains include CoV-2-s2m^Δ2-42^ which contains a deletion of the s2m element, CoV-2-s2m^2-5^ which contains four consecutive substitutions of the stem of the s2m, and CoV-2-s2m^2–5,39–42^ which contains complementary substitutions predicted to restore the secondary structure of the s2m. Infectious virus titer measured in plaque forming units per mL (PFU/mL) at 0, 24, 48, and 72 hours post-infection. A total of two experiments with three technical replicates each were conducted. No difference in the viral titer was detected by a two-way ANOVA with post-hoc testing by Dunnett’s multiple comparison test between WT and all mutants for Vero-hTMPRSS2 F(3,20)= 1.02, P= 0.40, and for Calu-3 cells F(3,20)= 0.48, P= 0.7). (**C**) Multi-step growth curve measuring the infectious viral titer using a clinical isolate of SARS-CoV-2 containing a partial deletion of the s2m element (SARS-CoV-2/WUSTL_000226_A131/2021), compared to a WA1-strain of SARS-CoV-2 with a D614G mutation. Viral titers were measured at 0, 24, 48, and 72 hours post-infection and no difference in titers was detected by a two-way ANOVA with post-hoc testing by Sidak’s multiple comparison test (F(1,20)= 2.34, P= 0.16). For all graphs, geometric means ± geometric standard deviations are depicted. Dotted line is the limit of detection of the assay.

### Growth of a clinical SARS-CoV-2 isolate with a deletion in the s2m

During genomic surveillance for SARS-CoV-2 variants in the St. Louis area, USA, we identified one genome (SARS-CoV-2/WUSTL_000226_A131/2021) that contained a deletion of 27 nucleotides that removes positions 22-43 of the s2m. This mutant is in the B1.2 lineage (Pango Lineages)^27^, contains the D614G variant in the spike protein, and has 99.83% nucleotide identity compared to the reference SARS-CoV-2 Wuhan genome (NC_045512.2). We were able to culture WUSTL_000226_A131/2021 and verified that the recovered virus maintained the deletion by sequencing. In a multi-step growth curve, we did not observe any significant difference in virus titer between WUSTL_000226_A131/2021 and a recombinant WA1-strain of SARS-CoV-2 engineered with D614G mutation at any timepoint (two-way ANOVA F[1,10]= 2.3, P= 0.16; **Fig. 4C**).

### The s2m element is dispensable for SARS-CoV-2 *in vivo*

We next determined if the SARS-CoV-2 s2m was important *in vivo* using the Syrian hamster model. Hamsters were intranasally inoculated with 1,000 PFU of WT or mutant viruses and weights were recorded daily for six days. Nasal washes and lungs were collected 3 and 6 days post infection. No difference in weight loss was observed between the hamsters inoculated with the WT and deletion or mutant s2m-containing viruses (mixed-effect model with Geisser-Greenhouse correction F[3,44]= 1.26, P= 0.30; **Fig. 5A**). No significant differences in infectious virus (lung tissue Kruskal-Wallis H(3)= 0.62, P= 0.89), viral RNA (lung tissue Kruskal-Wallis H(3)= 2.0, P= 0.57; nasal wash Kruskal-Wallis H(3)= 2.2, P= 0.54) were detected for CoV-2-s2m^Δ2-42^, CoV-2-s2m^2-5^, and CoV-2-s2m^2-5,39–42^ compared to WT (**Fig. 5B-D**). At six days post-inoculation, when antiviral immunes responses are expected to be activated, no difference in infectious virus was detected in the lungs of the mutant SARS-CoV-2 infected hamsters compared to WT (Kruskal-Wallis H(3)= 5.5 P= 0.14; **Fig. 5E**). The viral RNA load with CoV-2-s2m^Δ2-42^, CoV-2-s2m^2-5^, and CoV-2-s2m^2-5,39–42^ also showed no difference compared to WT from the lungs (Kruskal-Wallis H(3)= 6.2 P= 0.10) and nasal washes (Kruskal-Wallis H(3)= 2.8 P= 0.42; **Fig. 5F-G**). Combined, these data suggest that the original SARS-CoV-2 virus does not require the s2m for growth *in vitro* or *in vivo*.

**Figure 5:**
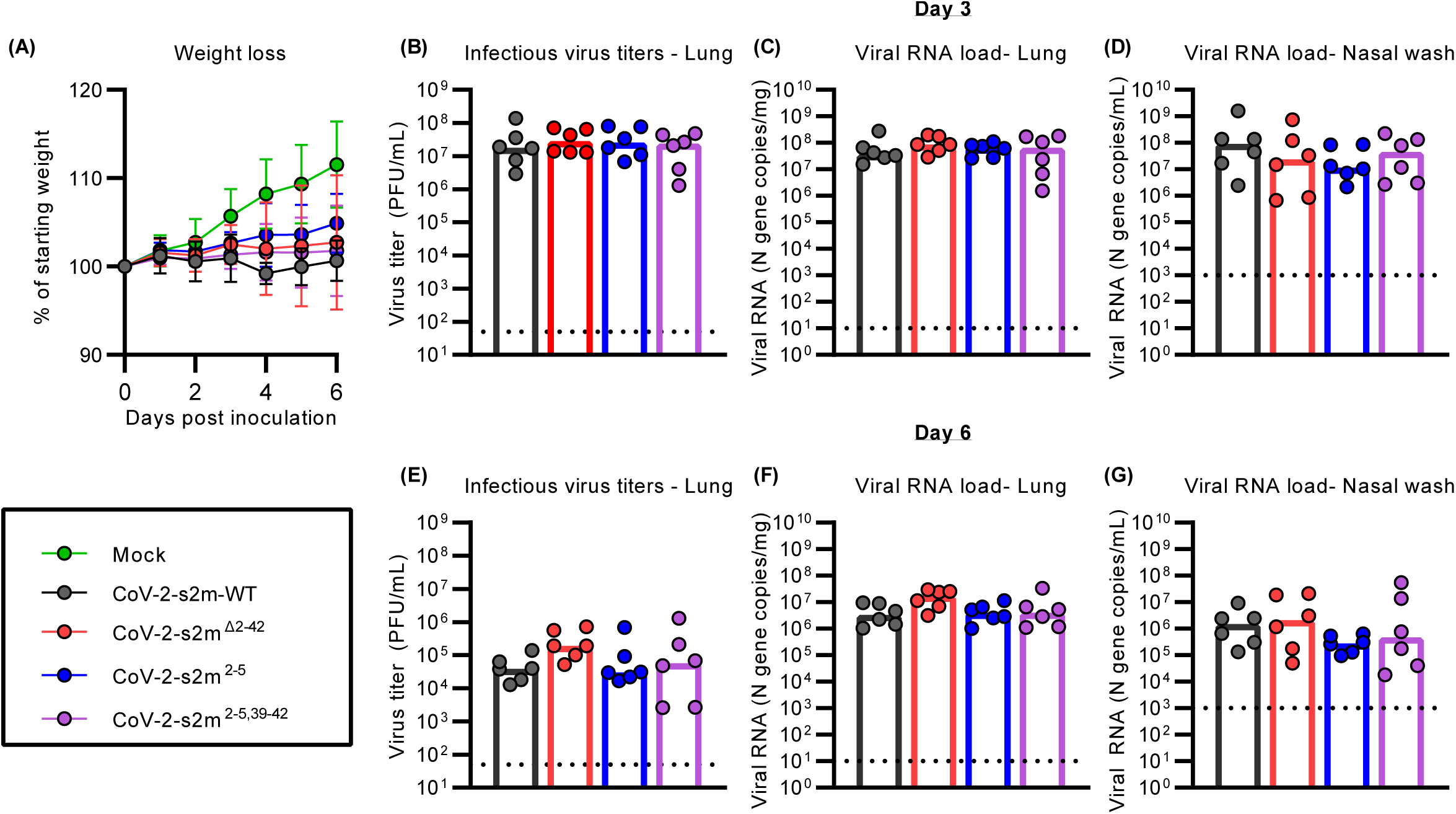
The s2m is dispensable for SARS-CoV-2 *in vivo.* Intranasal inoculation of hamsters with SARS-CoV-2 s2m WT and mutants including CoV-2-s2m^Δ2–42^ which contains a deletion of the s2m, CoV-2-s2m^2-5^ that contains four mutations of the stem of the s2m, and CoV-2-s2m^2-5,39–42^ that contains complementary mutations predicted to restore the secondary structure of the s2m. Hamsters were sacrificed at days 3 and 6. Two experiments were conducted, each experiment included three hamsters. (**A**) Mean hamster weight as percent of starting weight is graphed with error bars representing standard deviations. The weight of the hamsters infected with WT and mutant SARS-CoV-2 viruses were compared and no difference was detected using a mixed effect model with Geisser-Greenhouse correction (F[3,44] =1.26, P= 0.30). There was no difference in (**B**) Lung infectious viral titer (Kruskal-Wallis H(3)= 0.62 P= 0.89), (**C**) lung viral RNA (Kruskal-Wallis H(3)= 2.0 P= 0.57), and (**D**) nasal wash viral RNA (Kruskal-Wallis H(3)= 2.2 P= 0.54) from day 3 were identified comparing WT and s2m mutant viruses. At 6 days post inoculation, there was also no differences in (**E**) lung infectious viral titer (Kruskal-Wallis H(3)= 5.5 P= 0.14), (**F**) lung viral RNA (Kruskal-Wallis H(3)= 6.2 P= 0.10), and **(G)** nasal wash viral RNA (Kruskal-Wallis H(3)= 2.8 P= 0.42) comparing WT to mutant s2m viruses. For all graphs B-G, each dot is one animal. Dotted line represents the limit of detection of the respective assay.

## Discussion

Despite the first description of the s2m in 1997, no functional genetic studies on the role of the element in the virus lifecycle have been performed^1-6^. Here, we demonstrated that the s2m is critical for VA1 and important for HAstV1. In contrast, the deletion or mutation of the s2m in SARS-CoV-2 did not impact growth *in vitro* or *in vivo*. These results suggest that the importance and function of the s2m are virus-context dependent.

For VA1, we demonstrated the critical importance of G-C binding pairs in the s2m element, supporting the hypothesized secondary and tertiary structure. The pressure to maintain the structure of the s2m likely places constraints on the nucleotide diversity at each position (**Fig. 1A**). While position 18 can either be guanine or cytosine, position 24 is conserved with a complementary base that maintains a G-C binding pair. We demonstrated that VA1 tolerates both orientations of this G-C base pair without reduction of viral progeny (VA1-s2m-WT and VA1-s2m^18, 24^; **Fig. 2C**). Other G-C base pairs including positions 15-29, 16-28, and 17-25 are rigorously conserved across all s2m sequences (**Fig. 1A**). One challenge to testing these other G-C base pairs is that most astrovirus s2m elements partially overlap with the coding sequence of ORF2, limiting the mutations that can be introduced without affecting the amino acid sequence. We did find that VA1 tolerates disruption of the stop codon and the addition of 16 amino acids to the C-terminus of ORF2, as long as the s2m predicted structure was maintained (VA1-s2m^18, 24^ and VA1-s2m^12, 18, 24, 33^). ORF2 encodes the capsid protein, and it is known for HAstV1 and HAstV8 that the protein undergoes proteolytic cleavage^28–31^. It is possible that VA1 can tolerate the addition of 16 amino acids residues because this additional peptide is part of a larger peptide that is cleaved during the viral lifecycle.

We also identified that nucleotide positions that are not predicted to form G-C base pairs can be modified in the s2m (VA1-s2m^9^), but this resulted in reduced production of viral progeny (**Fig. 2C**). In the multiple-sequence alignment, adenine, cytosine, and guanine are observed to be present at this position (**Fig. 1A**). In the crystal structure for the SARS-CoV-1 s2m, position 9 was hypothesized to form a tertiary interaction with nucleotide 30 and may also bind a magnesium ion^4^. In future studies, we can determine if different nucleotide substitutions alter these interactions, providing further insights into what nucleotide substitutions can be tolerated by the s2m.

We corroborated the significance of the s2m for a different astrovirus, HAstV1, and the s2m was important but not critical. These findings highlight that the magnitude of the importance of the s2m is virus-dependent. Unlike VA1, HAstV1 may have greater tolerance to alterations to the s2m as the HAstV1 s2m tolerated replacement with the SARS-CoV-1 and SARS-CoV-2 s2m sequences in a replicon system^7^. Importantly, some astroviruses do not encode an s2m, including viruses of the MLB clade^10^. This observation is consistent with the finding that the s2m is not an integral lifecycle element for all astroviruses.

SARS-CoV-2 contains uracil at position 32 while all other representative s2m-containing viruses encode guanine (**Fig. 1A**). This position is predicted to form a G-C binding pair and we experimentally validated the significance of this genetic variant in VA1, as the recombinant VA1-s2m^32^ could not be rescued. This result is consistent with the significance of the G-C base pairs in maintaining the s2m structure. It is also consistent with the observation that the s2m element is dispensable in SARS-CoV-2. Other coronaviruses with an s2m, including SARS-CoV-1 and closely related bat coronaviruses RaTG13 (MN996532.2), BANAL-52 (MZ937000.1), and BANAL-236 (MZ937003.2) do not encode uracil at this position^32^. These findings suggest that the uracil genetic variant was a recent evolutionary event. Future identification of more closely related genomes to SARS-CoV-2 will provide further insights into the timing of the emergence of the uracil substitution.

Nonetheless, the presence of the uracil at position 32 in essentially all SARS-CoV-2 genomes (>99.99% on NCBI) raises major questions about the function of the SARS-CoV-2 s2m. Why does SARS-CoV-2 encode an s2m that contains a genetic variant in a position that is predicted to disrupt the secondary structure? Our results demonstrate the SARS-CoV-2 s2m is not critical for the viral lifecycle. There are multiple reports of mutations and deletions of the s2m in SARS-CoV-2 clinical isolates^23–26^, and the s2m region has, surprisingly, a higher mutational rate compared to other regions of the SARS-CoV-2 genome^23^. These observations, combined with our functional data raise the possibility that SARS-CoV-2 may have evolved an alternate mechanism to substitute for the function of the s2m. Alternatively, the s2m in SARS-CoV-2 may have a unique function but is still dispensable. Given the conservation of the uracil variant for SARS-CoV-2 across clinical isolates, it could provide an unknown fitness advantage.

One possibility for our findings is that the significance of the s2m is cell-line dependent. The s2m was important for both VA1 and HAstV1 in Caco-2 (human) cells, while the SARS-CoV-2 s2m was dispensable in Vero (African green monkey) or Calu-3 (human) cells, confirming the importance of the s2m is not specific to human cells. In addition, we noted different reductions of activity for the HAstV1 replicon system in HEK293T and Huh7.5 cells (**Fig 3C-D**). In the future, we can test if the s2m has different magnitudes of importance that dependent on expression levels of specific cellular pathways. In addition, there is currently no *in vivo* model in which to study either classic or VA-clade astroviruses. Future development of an animal model of astrovirus infection would enable us to determine if the observed *in vitro* phenotypes translate *in vivo*.

The mechanism behind the function of the s2m remains unclear. Various hypotheses have been proposed, but a clear function has not been established^2, 4^. Based on the presence of the s2m in viruses of five different viral families, it could act through a shared pro-viral mechanism. The s2m could be an important element that interacts with the host through s2m-host protein or RNA interactions. Any host pathway affected by the s2m is likely to be highly conserved since the s2m has been identified in viruses that are predicted to infect mammals, birds, and insects^6^. It is also possible that the s2m is a key element for viral RNA-RNA interactions during the viral lifecycle, as suggested by others^33, 34^.

In summary, we have defined differential roles for the s2m element that is virus-context dependent. Further studies will define the mechanistic basis as to why a highly conserved RNA element can lead to different functional roles in the viral lifecycle.

## Materials and Methods

### Cell culture conditions

BHK-21, HEK293T and Caco-2 cells were maintained in Dulbecco’s Modified Eagle Medium (DMEM) with L-glutamine (Gibco) supplemented with 10% fetal bovine serum (FBS) and 100 units/mL penicillin/streptomycin and incubated at 37°C and 5% CO_2_. Huh7.5.1 cells (obtained from Apath, Brooklyn, NY) were maintained in the same media supplemented with non-essential amino acids. Vero cells expressing human TMPRSS2 (Vero-hTMPRSS2) ^35^ or human ACE2 and human TMPRSS2 (Vero-hACE2-hTMPRSS2, gift from Drs. Graham and Creanga at NIH) and BSR cells (a clone of BHK-21) were cultured and maintained in DMEM supplemented with 5% FBS and 100 units/mL of penicillin and streptomycin. Vero-hTMPRSS2 and Vero-hACE2-hTMPRSS2 cells were maintained by selection with 5 µg/mL Blasticidin or 10 µg/mL puromycin respectively.

### Astrovirus VA1 reverse genetics system

The reference VA1 genome (NC_013060.1) was synthesized using gBlocks (Integrated DNA Technologies)^36^. This genome contains 4 or 5 nucleotide substitutions relative to previously propagated VA1 in cell culture, allowing for differentiation of recombinant virus^11, 12^. During construction of the genome, we identified significant bacterial toxicity of the VA1 sequence that hindered the assembly of the full genome in a plasmid. The toxicity localized to a region of ORF1a of the VA1 genome, so the genome was split into two plasmids (**Fig. S1A**). Plasmid p1629 contains a BamHI restriction site, T7 promoter (TAATACGACTCACTATAG) followed by the first 1629 nucleotides of the VA1 genome inserted into the pSMART plasmid (Lucigen). A second plasmid, p5023 contains a BamHI and BlpI restriction site, nucleotides 1610-6586 followed by a poly-A sequence of 18 adenosines, followed by EciI and SbfI restriction digest sites inserted into pUC19. To introduce mutations or deletions in the s2m of the VA1, synthetic DNA encoding for the nucleotides 6080-6586 were commercially synthesized with a poly-A tail added by overlap PCR and cloned into p5023 using unique enzyme restriction sites AgeI and SbfI (**Table S1**).

Due to low yields of p1629, the VA1 insert was amplified by PCR. Wild-type and mutant p5023 plasmids were linearized using BamHI and BlpI. NebBuilder HiFi assembly (New England Biolabs) was used to combine the fragments into the full-length VA1 sequence. The p1629 PCR product was combined with the p5023 fragment at a mass ratio of 1:2.5 using the manufacturer’s instructions by incubating the fragments for two hours at 50°C. The DNA was purified using a DNA column (Zymo Research). To cleave the 3′ end and generate a linear template for *in vitro* transcription, the VA1 DNA sequence was further digested overnight with EciI at 37°C. The DNA was again purified using a DNA column (Zymo Research) and the expected assembled genome DNA length was confirmed by gel electrophoresis. A total of 1 µg of DNA was used for *in vitro* transcription using mMESSAGE mMACHINE T7 Ultra Transcription kit (Invitrogen) with the 5′ end capped with anti-reverse cap analog (ARCA) and supplemented with GTP (1 µL per 20 µL reaction). After a two-hour incubation at 37°C, Turbo DNase was added and incubated for 15 minutes at 37°C. The RNA was purified using RNeasy (Qiagen) and quantified by Nanodrop (ThermoFisher). Two sets of *in vitro* transcribed RNA (IVT RNA) were generated for every VA1 mutant genome.

A total of 1.5 µg/well of VA1 IVT RNA was transfected into BHK-21 cells in 12-well plates using 3 µL/well of Lipofectamine MessengerMax (Invitrogen) and following the manufacturer’s protocol. The cells were incubated for 48 hours and then freeze-thawed three times with the lysate cleared by centrifugation (6,000 x g for 3 minutes). Using a previously established infection protocol for VA1, a total of 300 µL of cleared cell lysate was used for infection of Caco-2 cells in 6-well format ^37^. Five days after infection, the cells were freeze-thawed three times, and the cell lysate passaged three more times in Caco-2 cells to increase the viral titer and confirm maintenance of infectivity of each mutant genome. For all WT and mutant genomes, three separate experiments consisting of three independent replicates each were passaged in parallel, for a total of 9 replicates per condition. For all genomes that were viable, a 483 bp amplicon spanning the s2m region was amplified by RT-PCR and Sanger sequenced to confirm maintenance of the expected mutations.

A previously described fluorescent focus forming assay (FFA) was used to quantify the infectious titer of VA1 in focus forming units per mL (FFU/mL). Caco-2 cells were seeded into 96- well plates ^38^. Multi-step growth curves were completed using Caco-2 cells with samples collected at 1, 24, 48, and 96 hpi. A total of three separate experiments with three replicates each were conducted. The RNA from the cell fraction was extracted using a previously published protocol and viral RNA was quantified by qRT-PCR ^12, 37^. Combined cell lysate and supernatant were collected and used to quantify the infectious viral titer by FFA at 1, 48, and 96 hpi.

### Human astrovirus 1 reverse genetics system

Mutant s2m sequences were introduced using site-directed mutagenesis of a previously published plasmid encoding HAstV1 (pAVIC1) ^15^ and confirmed by sequencing. The resulting plasmids were linearized with XhoI prior to T7 RNA transcription using mMESSAGE mMACHINE T7 Transcription kit (Invitrogen). The pAVIC1- derived HAstV1 RNA was electroporated into BSR cells as previously described ^16^. For virus passaging and multistep growth curves, the collected supernatant was treated with 10 µg/mL

Type IX trypsin (Sigma) for 30 min at 37°C, diluted 5 times with serum-free media, and used for infection of Caco-2 cells. After three hours of incubation, the virus-containing media was replaced with serum free media containing 0.6 µg/mL trypsin, and cells were incubated for 48-72 hours until the appearance of cytopathic effect. After 3 freeze-thaw cycles, viral stocks were titrated using Caco-2 cells on 96-well plates in the absence of trypsin, enabling single-round infection ^16^. The immunofluorescence-based detection with 0.26 µg/mL of 8E7 astrovirus antibody (Santa Cruz Biotechnolgy, sc-53559) was combined with infrared detection readout and automated LI-COR software-based quantification. The titers of HAstV1 are determined as infectious units per ml (IU/mL). Both passaging and multi-step growth curves were performed using an MOI of 0.1. For multi-step growth curves, individual infections were collected at 1, 6, 12, 24, 36, 48, 72, and 96 hpi and saved for virus quantification, with a total of three separate experiments.

### Human astrovirus 1 replicon system

A previously published HAstV1 replicon system was utilized, based on the pAVIC1 plasmid where the subgenomic RNA encodes the foot-and-mouth disease virus 2A sequence followed by a Renilla luciferase (RLuc) reporter sequence ^16^. The s2m deletion was introduced using available restriction sites and confirmed by sequencing. The resulting plasmids were linearized with XhoI and IVT was produced using mMESSAGE mMACHINE T7 Transcription kit, purified using Zymo RNA Clean & Concentrator kit and quantified by nanodrop. Huh7.5.1 and HEK293T cells were transfected in triplicate with HAstV1 replicon IVT RNA using Lipofectamine 2000 (Invitrogen), following a previously described protocol in which suspended cells are added directly to the RNA complexes in 96-well plates ^7^. The transfected cells were incubated for 18 hours. Replicon activity was calculated as the ratio of Renilla (subgenomic reporter) to Firefly (co-transfected loading control RNA, cap-dependent translation) using Dual-Luciferase Stop & Glo Reporter Assay System (Promega) and normalized by the same ratio for the control WT replicon. Three independent experiments, each in triplicate, were performed to confirm the reproducibility of results.

### SARS-CoV-2 reverse genetics system

All work with potentially infectious SARS-CoV-2 particles was conducted under enhanced biosafety level 3 (BSL-3) conditions and approved by the institutional biosafety committee of Washington University in St. Louis. The prototypic SARS-CoV-2 Wuhan genome (NC_045512.2) was split into 7 fragments (**Fig S1C**), named A to G, and each DNA fragment was commercially synthesized (GenScript). A T7 promoter sequence was introduced at the 5′ end of fragment A, and a poly-A sequence of 22 adenosines was introduced at the 3′ end of fragment G. In addition, NotI and SpeI sites were introduced at the 5′ end of fragment A before the T7 promoter and the 3′ end of fragment G after the poly-A sequence, respectively. To ensure seamless assembly of the full virus DNA genome, the 3′ end of fragment A, both ends of fragments B-F and the 5′ end of fragment G were appended by class II restriction enzyme recognition sites (BsmBI and BsaI) (**Fig S3C**). Fragments A and C-G were cloned into plasmid pUC57 vector and amplified in *E. Coli* DH5α strain. The bacteria toxic fragment B was cloned into low copy inducible BAC vector pCCI and amplified through plasmid induction in EPI300. Low copy plasmids were extracted by NucleoBond Xtra Midi kit (MACHEREY-NAGEL) with the other plasmids were extracted by plasmids midi kit (QIAGEN) according to the manufacturers’ protocols. To generate mutant s2m sequences, the SARS-CoV-2 s2m sequence in the G fragment was mutated by site-directed mutagenesis (**Table S1**). Mutations and deletions in the s2m element were confirmed by Sanger sequencing. To assemble the full-length SARS-CoV-2 genome, each DNA fragment in plasmid was digested by corresponding restriction enzymes, DNA fragments were recovered by gel purification columns (New England Biolabs), and the seven fragments were ligated at equal molar ratios with 10,000 units of T4 ligase (New England Biolabs) in 100 µL at 16°C overnight. The final ligation product was incubated with proteinase K in the presence of 10% SDS for 30 minutes, extracted twice with equal volume of phenol:chloroform:isoamyl alcohol (25:24:1; ThermoFisher), isopropanol precipitated, air dried and resuspended in RNase/DNase free water. The DNA was analyzed on a 0.6% agarose gel. Full length genomic SARS-CoV-2 RNA was *in vitro* transcribed using mMESSAGE mMACHINE T7 Ultra Transcription kit (Invitrogen) following the manufacturer’s protocol. Four µg of DNA template was added to the reaction mixture, supplemented with GTP (7.5 µL per 50 μL reaction). *In vitro* transcription was done overnight at 32°C. Afterwards, the template DNA was removed by digestion with Turbo DNase for 30 minutes at 37°C. The *in vitro* transcript (IVT RNA) mixture was used directly for electroporation. To further enhance rescue of recombinant virus, we also generated SARS-CoV-2 nucleocapsid (N) gene RNA. The SARS-CoV-2 *N* gene was PCR amplified from plasmids pUC57-SARS-CoV-2-N (GenScript) using forward primers with T7 promoter and reverse primers with poly(T)34 sequences. The *N* gene PCR product was gel purified and used as the template for *in vitro* transcription using the same mMESSAGE mMACHINE T7 Transcription Kit with 1 µg of DNA template, 1 µL of supplemental GTP in a 20 µL reaction volume. For SARS-CoV-2 IVT RNA electroporation, low passage BHK-21 cells were trypsinized and resuspended in cold PBS as 0.5×10^7^ cells/mL. A total of 20 μg of SARS-CoV-2 IVT RNA and 20 μg of *N* gene *in vitro* transcript were added to resuspended BHK-21 cells in a 2 mm gap cuvettes and electroporated with setting at 850 V, 25 μF, and infinite resistance for three times with about 5 seconds interval in between pulses. The electroporated cells were allowed to rest for 10 minutes at room temperature and were then co-cultured with Vero-hACE2-hTMPRSS2 cell at a 1:1 ratio in a T75 culture flask. Cell culture medium was changed to DMEM with 2% FBS the next day. Cytopathic effect (CPE) was monitored for five days, and cell culture supernatant were harvested for virus titration.

### SARS-CoV-2 Syrian hamster infection model

Animal studies were carried out in accordance with the recommendations in the Guide for the Care and Use of Laboratory Animals of the National Institutes of Health. The protocols were approved by the Institutional Animal Care and Use Committee at the Washington University School of Medicine (assurance number A3381– 01). Five to six-week-old male hamsters were obtained from Charles River Laboratories and housed at Washington University. Next, the animals were challenged via intranasal route with 1,000 PFU of the recombinant WT or s2m mutant SARS-CoV-2 viruses under enhanced BSL-3 conditions^39^. Animal weights were measured daily for the duration of the experiment. Three and six days after the inoculation, the animals were sacrificed, and lung tissues and nasal washes were collected. The nasal wash was performed with 1.0 mL of PBS containing 0.075% BSA, clarified by centrifugation for 10 minutes at 2,000 × g and stored at -80°C. The left lung lobe was homogenized in 1.0 mL DMEM, clarified by centrifugation (1,000 × g for 5 minutes) and used for viral titer analysis by plaque assay and RT-qPCR using primers and probes targeting the *N* gene. For viral RNA quantification, RNA was extracted using RNA isolation kit (Omega Bio-tek). SARS-CoV-2 RNA levels were measured by one-step quantitative reverse transcriptase PCR (RT-qPCR) TaqMan assay as described previously using a SARS-CoV-2 nucleocapsid (N) specific primers/probe set from the Centers for Disease Control and Prevention (F primer: GACCCCAAAATCAGCGAAAT; R primer: TCTGGTTACTGCCAGTTGAATCTG; probe: 5′-FAM/ACCCCGCATTACGTTTGGTGGACC/3′-ZEN/IBFQ)^40^. Viral RNA was expressed as (N) gene copy numbers per mg for lung tissue homogenates or mL for nasal swabs, based on a standard included in the assay, which was created via *in vitro* transcription of a synthetic DNA molecule containing the target region of the N gene.

### SARS CoV-2 growth curves and titration assays

Vero-hTMPRSS2 and Calu-3 cells were grown to confluency. Cells were inoculated with a multiplicity of infection (MOI) of 0.001 of recombinant WT or mutant SARS-CoV-2 and culture supernatant was collected at 1, 24, 48, and 72 hpi and saved for viral quantification by plaque assay on Vero-hACE2-hTMPRSS2 cells in 24-well plates as described ^41^. A total of two separate experiments with three replicates each were conducted.

### Propagation of a clinical isolate of SARS-CoV-2 containing a deletion of the s2m

As part of ongoing SARS-CoV-2 variant surveillance, a random set of RT-PCR positive respiratory secretions from the Barnes Jewish Hospital Clinical microbiology laboratory were subjected to whole genome sequencing using the ARTIC primer amplicon strategy ^42^. This study was approved by the Washington University Human Research Protection Office (#202004259). From the sequences generated, one genome (SARS-CoV-2/human/USA/WUSTL_000226_A131/2021; Genbank # OM831956) had a 27-nucleotide deletion that removed 22 nucleotides from the 3′ end of the s2m element (**Fig S4**). The spike protein of this virus harbored L18F, D614G, and E780Q mutations, demonstrating that this virus belonged the original B.1 lineage of SARS-CoV-2. This virus was expanded twice on Vero-hACE2-hTMPRSS2 cells and the virus titer was determined by plaque assay. The P2 of WUSTL_000226_A131/2021 was sequenced by NGS to confirm the presence of the s2m deletion and rule out any tissue culture adaptations in the rest of the genome.

### Sequencing of reverse genetics rescued VA1, HAstV1, and SARS-CoV-2 virus

For VA1, 100 µL of whole cell lysate was used for RNA column-based extraction (Zymo). Ribosomal RNA was removed from total RNA by Ribo-Zero depletion (Illumina) and libraries constructed using Stranded Total RNA Prep kit (Illumina). Libraries were pooled and sequenced using an Illumina NextSeq system. Sequencing data was processed through IDseq/Chan Zuckerberg ID^43^, and consensus genomes were generated with single nucleotide polymorphisms identified by the default settings.

For SARS-CoV-2, 200 µL of supernatant containing recombinant SARS-CoV-2 was collected and added to 600 µL TRK lysis buffer plus beta-mecaptoethanol (BME) according to the E.Z.N.A. Total RNA Kit I (Omega Bio-tek). Total RNA was extracted according to the manufacturer’s protocol and eluted into RNase/DNase free H_2_O and used for library construction. Ribosomal RNA was removed from total RNA by Ribo-Zero depletion (Illumina). Indexed sequencing libraries were prepared using TruSeq RNA library preparation kit (Illumina), pooled and then sequenced using Illumina NextSeq system. The sequencing data were analyzed using the LoFreq pipeline to call the mutations in the entire virus genome ^44^. In brief, Illumina sequencing data were filtered using fastp (https://github.com/OpenGene/fastp) trimming bases with a Phred quality score ≥Q30. High-quality reads were then aligned to the SARS-CoV-2 reference genome sequence (NC_045512.2) using BWA ^45^. SAMtools was used to sort, index and remove duplicates from bam files, and local realignment and variant calling were achieved by LoFreq to generate the mutant report file ^46^.

For HAstV1, the virus RNA was isolated from the virus stocks using Direct-zol RNA MicroPrep (Zymo research) and used for RT-PCR and Sanger sequencing of the entire virus genome.

### Multiple sequence alignment

Representative s2m sequences were aligned in MegaX using Muscle ^47^. Sequences were visualized using Jalview 2 ^48^. Representative sequences included: Astrovirus VA1 (NC_013060.1), Astrovirus VA2 (GQ502193.2), Human astrovirus 1 (NC_001943.1), Murine astrovirus (NC_018702.1), SARS-CoV-1 (NC_004718.3), SARS-CoV-2 (NC_045512.2), Bat coronavirus RaTG13 (MN996532.2), Bat coronavirus BANAL-52 (MZ937000.1), Bat coronavirus BANAL-236 (MZ937003.2), Avian infectious bronchitis virus (NC_001451.1), Thrush coronavirus HKU12-600 (NC_011549.1), Hubei tetragnatha maxillosa virus 9 (KX884602.1), Equine rhinitis B virus 1 (NC_003983.1), Bat picornavirus 3 (NC_015934.1), Dog norovirus (FJ692500.1).

### RNAfold analysis of s2m sequences and mutants

The secondary structures of the s2m sequences from SARS-CoV-1, SARS-CoV-2, HAstV1, and VA1 were predicted using the RNAfold Server ^49^ using default settings with the 2004 Turner model of RNA parameters and no dangling end energies ^50^. Mutant s2m sequences were also evaluated using this program. The minimum free energy of the predicted secondary structure was calculated using RNAfold.

### Analysis of SARS-CoV-2 s2m sequences using the NCBI database

We downloaded SARS-CoV-2 complete genome sequences from NCBI which were deposited before November, 2021 as a single FASTA file. The FASTA dataset was processed with a published SARS-CoV-2- freebayes pipeline to call all the variants along the genome ^51^. In brief, the single SARS-CoV-2 complete genome FASTA file was decomposed into individual FASTA genome sequences. Then, each FASTA genome was aligned individually against the SARS-CoV-2 reference sequence (NC_045512.2) using Minimap2 ^52^. Variant calling was performed on each BAM file using Freebayes variant caller to produce the VCF files. Variants in the s2m position 32 for all the VCF files were extracted and reported.

### Statistical analysis

Data was graphed using Prism 9.3.1 (GraphPad). Parametric and non-parametric comparisons were made where appropriate. Post hoc testing of Kruskal Wallis comparisons was completed with correction of multiple comparisons by Dunn’s multiple test correction. For multi-step growth curves, mutants were compared to WT by logarithmically transforming the data and comparing using a two-way ANOVA in Prism after excluding the one hour timepoint. Post hoc testing was conducted using either Sidak’s multiple comparison testing or hamster weight data analyzed by mixed-effect model with Geisser-Greenhouse correction in Prism. Adjusted P values ≤ 0.05 were considered significant.

## AUTHOR CONTRIBUTIONS

A.B.J. and M.C.O. performed experiments with astrovirus VA1. H.J., C.F., T.L.B., T.L.D. and H. H. H. performed experiments with SARS-CoV-2. T.L.B., and T.L.D. performed the animal experiments. K.S. performed viral load analysis by RT-qPCR. T.L.B. and T.L.D. performed viral load analysis by plaque assay. S.T., A.J., B.F., J.A.B., S.A.H., and B.A.P. analyzed sequencing data. V.L. conducted the experiments with HAstV1, wrote corresponding parts of the manuscript. D.W., A.B.J., A.C.M.B., and V.L provided supervision and acquired funding. A.B.J. summarized the data and performed the statistical analysis. A.B.J., H.J., D.W., and A.C.M.B. wrote the initial draft, with the other authors providing editorial comments.

---

**Supplemental Table 1:**
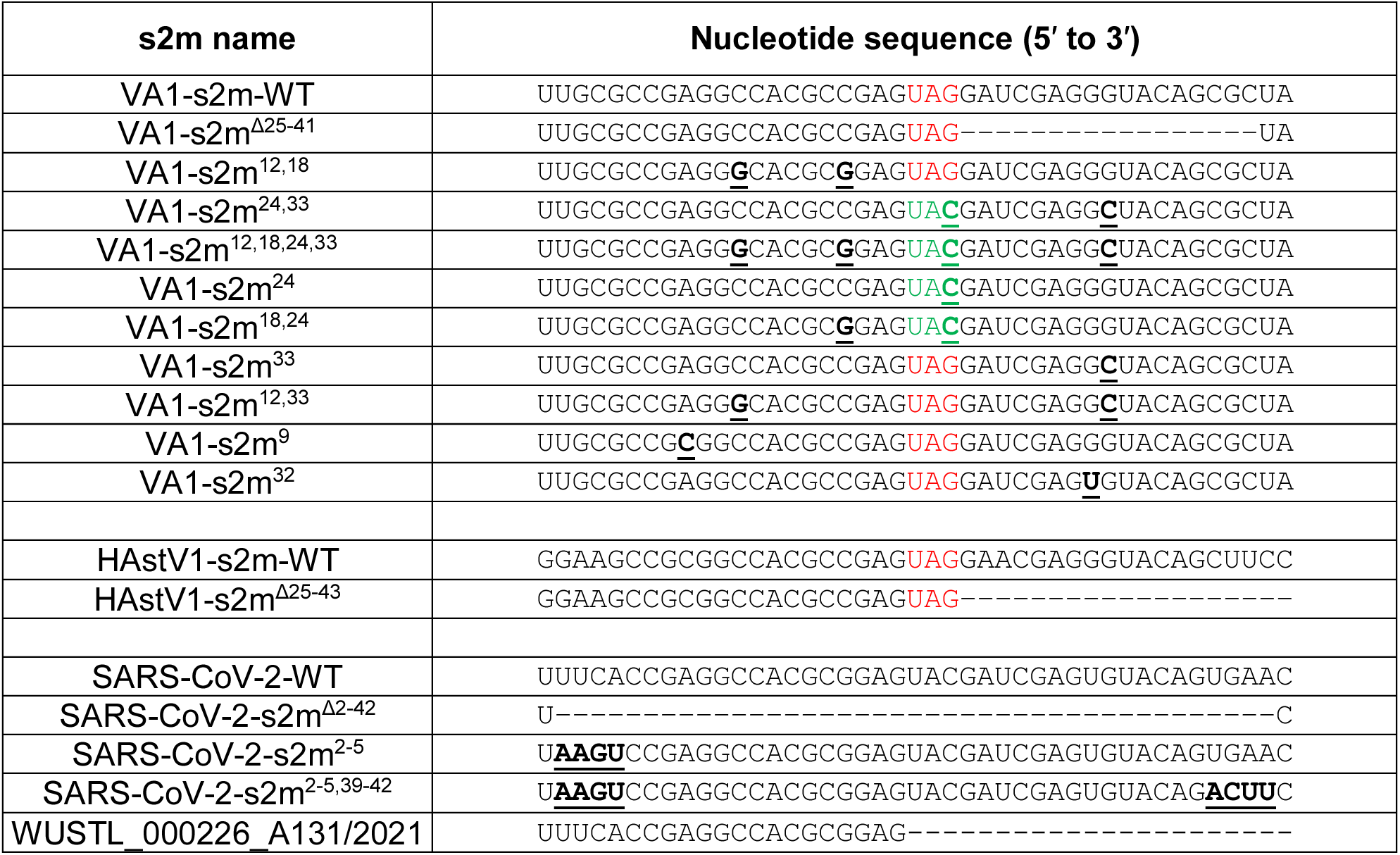
Nucleotide sequences of the different s2m sequences used in this study. Mutations relative to the wild-type virus s2m are in bold and underlined. Deleted nucleotides represented by a dash. Predicted stop codon for ORF2 for astroviruses highlighted in red, mutated stop codon sequence highlighted in green. WUSTL_000226_A131/2021 is a clinical isolate of SARS-CoV-2 that contains a deletion of 27 nucleotides, which includes 22 nucleotides of the s2m

**Supplemental Table 2:**
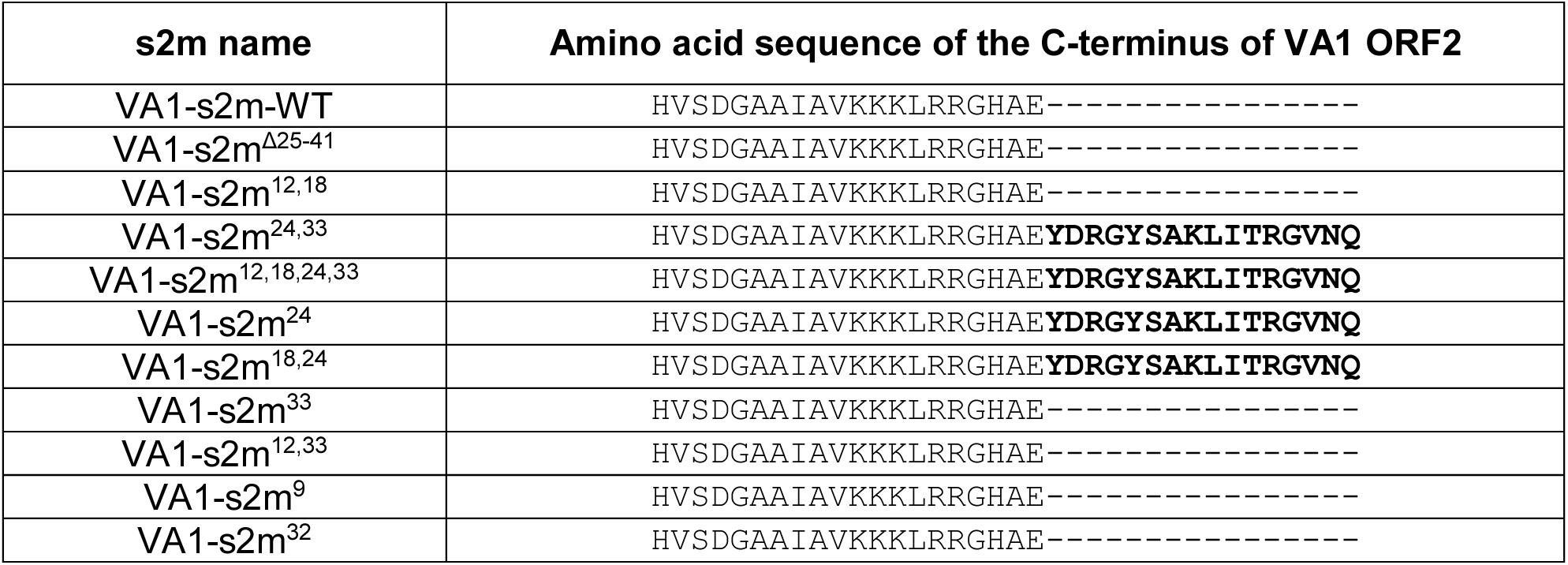
Amino acid sequences of VA1 wild-type and s2m mutant viruses. Depicted are the C-terminus amino acid sequence for VA1 ORF2 of the recombinant viruses. In VA1-s2m-WT and most other s2m mutant viruses, the predicted ORF2 gene contains a stop codon after the glutamic acid (E) residue. Mutants VA1-s2m^24^, VA1-s2m^18, 24^, VA1-s2m^24, 33^, and VA1-s2m^12, 18, 24, 33^ contain a mutation (position 24 of the s2m) that disrupts the stop codon resulting in addition of 16 amino acids to C-terminal end of ORF2 (bold).

| s2m name | Amino acid sequence of the C-terminus of VA1 ORF2 |
| --- | --- |
| VA1-s2m-WT | HVSDGAAI AVKKKLRRGHAE----- |
| VA1-s2m<sup>Δ25-41</sup> | HVSDGAAI AVKKKLRRGHAE----- |
| VA1-s2m<sup>12,18</sup> | HVSDGAAI AVKKKLRRGHAE----- |
| VA1-s2m<sup>24,33</sup> | HVSDGAAI AVKKKLRRGHAE**YDRGYSAKLITRGVNQ** |
| VA1-s2m<sup>12,18,24,33</sup> | HVSDGAAI AVKKKLRRGHAE**YDRGYSAKLITRGVNQ** |
| VA1-s2m<sup>24</sup> | HVSDGAAI AVKKKLRRGHAE**YDRGYSAKLITRGVNQ** |
| VA1-s2m<sup>18,24</sup> | HVSDGAAI AVKKKLRRGHAE**YDRGYSAKLITRGVNQ** |
| VA1-s2m<sup>33</sup> | HVSDGAAI AVKKKLRRGHAE----- |
| VA1-s2m<sup>12,33</sup> | HVSDGAAI AVKKKLRRGHAE----- |
| VA1-s2m<sup>9</sup> | HVSDGAAI AVKKKLRRGHAE----- |
| VA1-s2m<sup>32</sup> | HVSDGAAI AVKKKLRRGHAE----- |

**Supplemental Figure 1:**
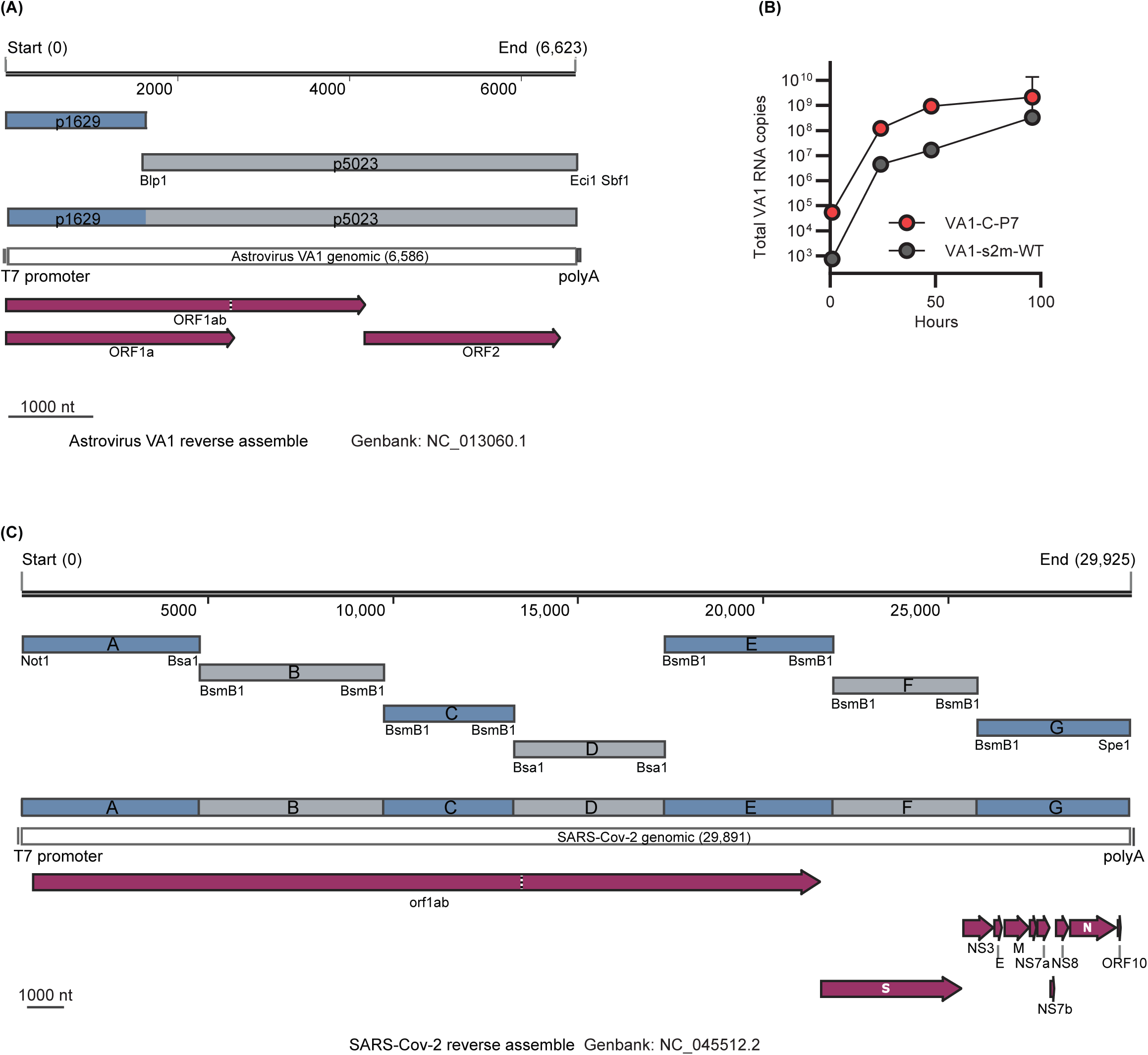
Design of the Astrovirus VA1 and SARS-CoV-2 reverse genetics. (**A**) Schematic of the Astrovirus VA1 full genome and fragment construction for the DNA genome assembly. T7 promoter, polyA sequence, Astrovirus VA1 encoded ORFs were indicated on the genome map in purple. Restriction sites were shown in its position along the genome on each fragment. (**B**) Multi-step growth curve with VA1 comparing an isolate that has been previously propagated in Caco-2 cells for seven passages (VA1-C-P7) and contains 5 mutations relative to the reference VA1 genome (NC_013060.1). VA1-s2m-WT is a recombinant virus generated from the VA1 reverse genetics system. RNA was measured at 0, 24, 48, and 96 hours post-infection. Error bars represent geometric standard deviation. (**C**) Schematic of the SARS-CoV-2 full genome and fragment construction for the cDNA genome assembly. T7 promoter, polyA sequence, SARS-CoV-2 encoded ORFs were indicated on the genome map in purple. Restriction sites were shown in its position along the genome on each fragment.

**Supplemental Figure 2:**
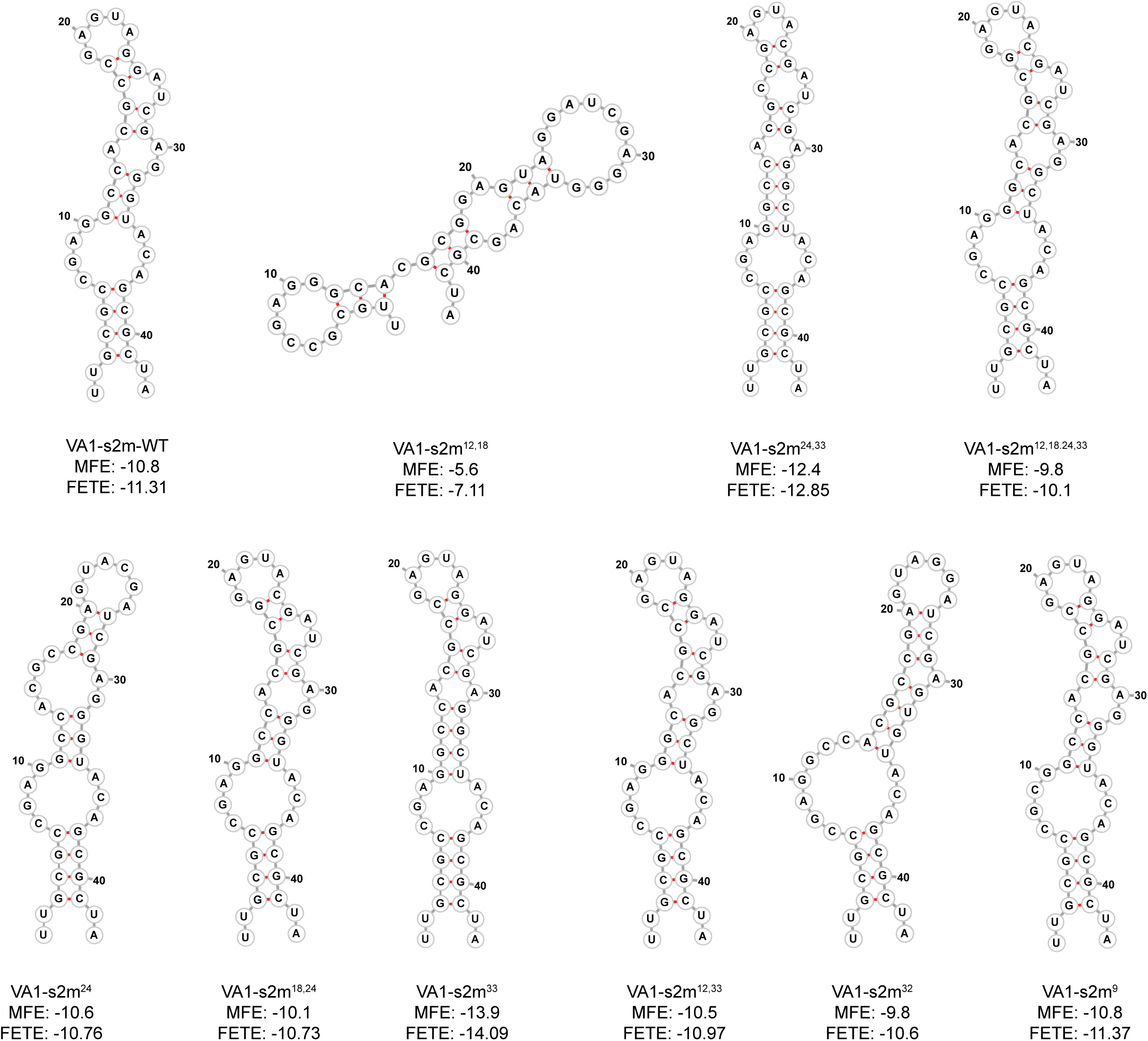
Predicted VA1 WT and mutant s2m secondary structures. The WT and mutant s2m RNA sequences substituted into the VA1 genome were analyzed using RNAfold. The predicted minimum free energy secondary structure is depicted including calculated the minimum free energy (MFE) and free energy of the thermodynamic enemble (FETE).

**Supplementary Figure 3:**
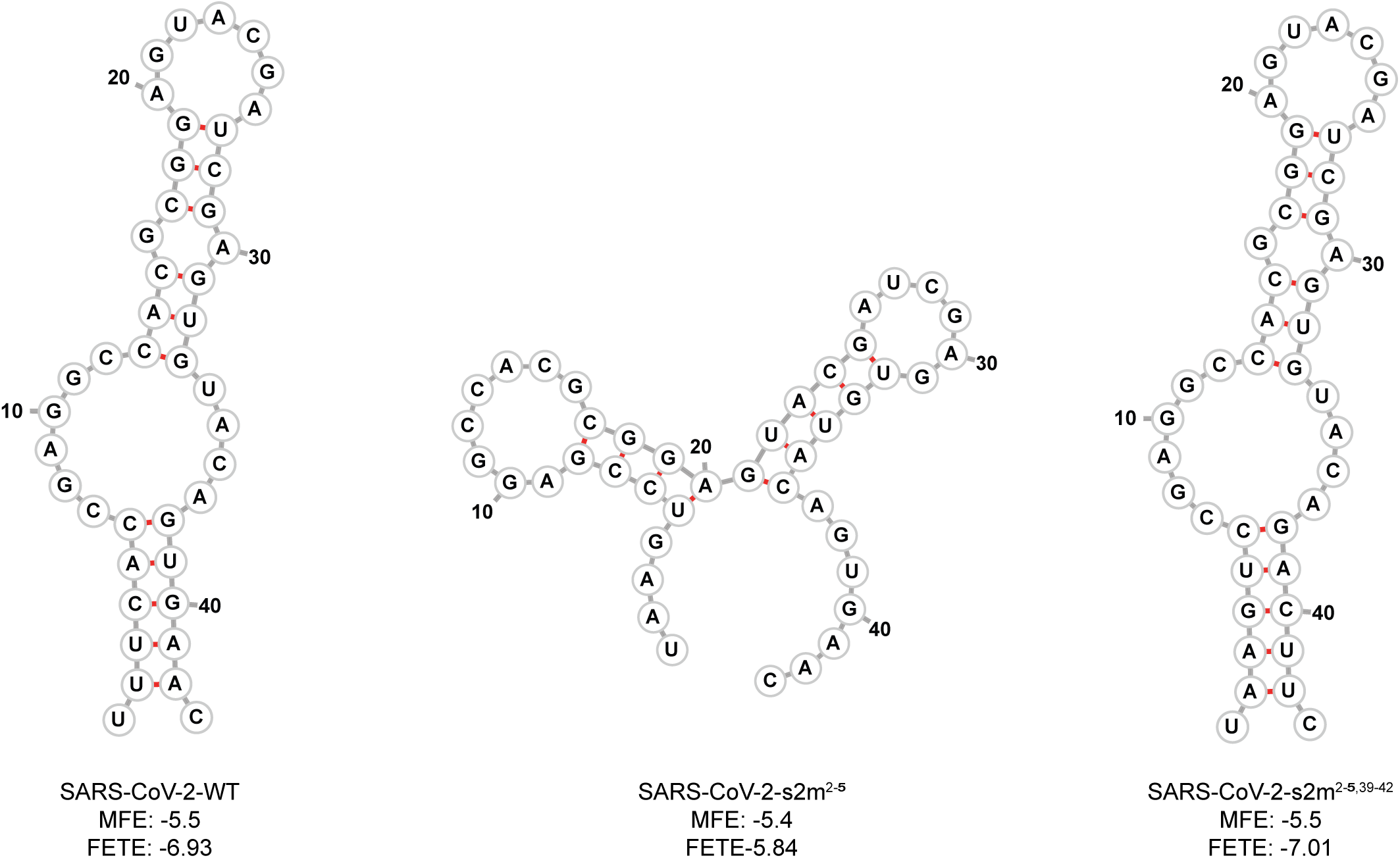
Predicted SARS-CoV-2 WT and mutant s2m secondary structures. Using RNAfold, the predicted minimum free energy secondary structure of the WT and mutant s2m RNA sequences substituted into the SARS-CoV-2 genome are shown. The calculated the minimum free energy (MFE) and free energy of the thermodynamic enemble (FETE) of each sequence is also shown.

